# The serogroup B meningococcal outer membrane vesicle-based vaccine 4CMenB induces cross-species protection against *Neisseria gonorrhoeae*

**DOI:** 10.1101/2020.05.13.094177

**Authors:** Isabelle Leduc, Kristie L. Connolly, Afrin Begum, Knashka Underwood, Nazia Rahman, Stephen Darnell, Jacqueline T. Balthazar, William M. Shafer, Andrew N. Macintyre, Gregory D. Sempowski, Ann E. Jerse

## Abstract

There is a pressing need for a gonorrhea vaccine due to the high disease burden associated with gonococcal infections globally and the rapid evolution of antibiotic resistance in *Neisseria gonorrhoeae* (*Ng*). Current gonorrhea vaccine research is in the stages of antigen discovery and the identification of protective immune responses, and no vaccine has been tested in clinical trials in over 30 years. Recently, however, it was reported in a retrospective case-control study that vaccination of humans with a serogroup B *Neisseria meningitidis* (*Nm*) outer membrane vesicle (OMV) vaccine (MeNZB) was associated with reduced rates of gonorrhea. Here we directly tested the hypothesis that *Nm* OMVs induce cross-protection against gonorrhea in a well-characterized female mouse model of *Ng* genital tract infection. We found that immunization with the licensed *Nm* OMV-based vaccine 4CMenB (Bexsero®) significantly accelerated clearance and reduced the *Ng* bacterial burden compared to administration of alum or PBS. High titers of serum IgG1 and IgG2a and vaginal IgG1 that cross-reacted with *Ng* OMVs were induced by vaccination via either the subcutaneous or intraperitoneal routes, and a 4-fold increase in the serum bactericidal_50_ titers was detected against the challenge strain. Antibodies from vaccinated mice recognized several surface proteins in a diverse collection of *Ng* strains, including PilQ, BamA, MtrE, PorB, and Opa, and 4CMenB-induced antibodies bound PilQ and MtrE in native form on the surface of viable bacteria. In contrast, the antibodies were only cross-reactive against lipooligosaccharide species from a few *Ng* strains. Our findings directly support epidemiological evidence that *Nm* OMVs confer cross-species protection against *Ng* and implicate several *Ng* surface antigens as potentially protective targets. This work also validates the murine infection model as a relevant experimental system for investigating mechanisms of vaccine-mediated protection against gonorrhea.

**Author summary:** Over 78 million *Neisseria gonorrhoeae (Ng)* infections occur globally each year and control of gonorrhea through vaccination is challenged by a lack of strong evidence that immunity to gonorrhea is possible. This contention was recently challenged by epidemiological evidence suggesting that an outer membrane vesicle (OMV) vaccine from the related species *Neisseria meningitidis* (*Nm*) protected humans against gonorrhea. Here we provide experimental evidence in support of this hypothesis by demonstrating that a licensed, modified version of this *Nm* OMV-based vaccine accelerates clearance of *Ng* in a mouse infection model. These results confirm the possibility cross-species protection and are important in that they support the biological feasibility of vaccine-induced immunity against gonorrhea. We also showed that several *Ng* outer membrane proteins are recognized by antisera from vaccinated mice that may be protective targets of the vaccine. Additionally, our demonstration that a vaccine that may reduce the risk of gonorrhea in humans protects mice against *Ng*, a highly host-restricted pathogen, validates the mouse model as a potentially useful tool for examining mechanisms of protection, which could be exploited in the development of other candidate gonorrhea vaccines.

## Introduction

An estimated 78 million new gonorrheal infections occur each year worldwide [1] and rates are rising globally, with a 67% increase in reported infections in the U.S. between 2014 and 2018 [2]. Caused by the Gram-negative bacterium *Neisseria gonorrhoeae (Ng)*, gonorrhea is associated with significant morbidity and mortality that disproportionately affects women and newborns. Ascending lower urogenital tract infection can occur in both sexes to cause epididymis, endometritis and salpingitis, but is more frequent in females. *Ng* pelvic inflammatory disease can be asymptomatic or acute, and is associated with ectopic pregnancy, infertility and chronic pelvic pain. Disseminated gonococcal infection can occur in either gender [3]. Transmission of gonorrhea to neonates from infected mothers can cause acute neonatal conjunctivitis [4] and there is a clear association between maternal gonorrhea, low-birth weight and pre-mature delivery [5]. The impact of gonorrhea on human health is amplified by its role in increasing both transmission and susceptibility to the human immunodeficiency virus (HIV) [6, 7].

Gonorrhea is classified as an urgent public health threat due to decreasing susceptibility to the last remaining reliable monotherapy for gonorrhea, the extended-spectrum cephalosporins. Dual therapy with high-dose ceftriaxone and azithromycin is currently recommended for empirical treatment of gonorrhea in many countries. However, *Ng* susceptibility to these antibiotics continues to decrease world-wide [8], and alarmingly, treatment failures due to strains that are resistant to these antibiotics have been reported [9, 10]. New antibiotics are under development [11, 12]; however, the call for a gonorrhea vaccine has been reinvigorated by the evolutionary success of the gonococcus in outrunning public health efforts to contain it through antibiotic therapy [13, 14].

Early vaccine research was challenged by the discovery that several *Ng* surface molecules are phase or antigenically variable. There was also no animal model other than chimpanzees for analyzing host responses and systematic testing of immunogens, and two published clinical trials using a killed whole cell vaccine [15] or purified pili [16] were unsuccessful despite earlier small in-house studies that showed protection from urethral challenge in human male volunteers [17]. Since this time, several conserved and semi-conserved vaccine antigens that elicit bactericidal antibodies or inhibit target function have been identified, some of which show protection in a well-characterized mouse genital tract infection model [13]. How well the mouse model predicts vaccine efficacy in humans is not known, however, due to the strict host-specificity of *Ng* and a lack of information on correlates of immune protection in humans. There is little immunity to natural infection in humans and mice [18], and there is growing evidence that the adaptive response to *Ng* infection is suppressed. As recently reviewed by Lovett and Duncan [19], human humoral immune responses to *Ng* to infection are modest at best and the analysis thereof is complicated by pre-existing antibodies to carbohydrate and protein surface antigens that are induced by commensal *Neisseria sp*., although antibodies to some antigens are increased by infection [20]. Human cellular responses to *Ng* infection are less well studied, but appear to be driven by a Th17 pro-inflammatory response. Th1 responses, in contrast, appear suppressed [20], and several pathways that result in reduced antigen presentation and, or inhibition of T cell responses to *Ng* have been identified using human immune cells and experimentally infected mice [21-25].

Recent epidemiological evidence, however, suggests immunity to gonorrhea can be achieved in humans through vaccination with outer membrane vesicles (OMVs) of the related species, *Neisseria meningitidis* (*Nm*). In this cross-sectional study, vaccination of individuals with the serogroup B meningococcal vaccine MeNZB, which consisted of OMVs from an endemic New Zealand strain, was associated with a reduced rate of gonorrhea in adolescents and adults aged 15-30 years old [26]. Using cases of chlamydia as a control, the estimated effectiveness of this meningococcal vaccine against gonorrhea was predicted to be 31%. These data are the first controlled evidence in humans in over 40 years that vaccine-induced protection against gonorrhoea is possible. A similar finding was suggested by epidemiological studies on *Nm* OMV vaccines in Cuba and Norway [27].

To directly test the hypothesis that *Nm* OMV-based vaccines induce cross-species protection against *Ng*, here we evaluated the *in vivo* efficacy of the licensed 4CMenB (“4 Component Meningitis B”; Bexsero^®^) vaccine in a female mouse model of *Ng* lower genital tract infection. 4CMenB consists of *Nm* OMVs from the *Nm* strain used in the MenNZB vaccine and five recombinant *Nm* proteins [28], only one of which, the neisserial heparin-binding antigen (NHBP), is a feasible vaccine target for gonorrhea [29, 30]. Our results show that 4CMenB significantly reduces the *Ng* bioburden, accelerates clearance of infection, and induces antibodies that recognize several *Ng* proteins, at least three of which are promising vaccine targets. These findings are consistent with epidemiological data that suggest cross-species protection against gonorrhea is possible and validate the gonorrhea mouse model as a useful experimental system for studying vaccine-mediated correlates of protection against this human disease.

## Results

### Optimization of the immunization regimen to induce serum and vaginal antibodies

The recommended dosing regimen for 4CMenB in humans is two 500 µL doses given intramuscularly, four weeks apart. As a preliminary step for mouse immunization/challenge studies, we immunized BALB/c mice with 20, 125, or 250 µL of the 4CMenB vaccine on days 1 and 28 by the subcutaneous (SC) or intraperitoneal (IP) routes to assess safety and immunogenicity. A dose-response in serum IgG1 titers against the 4CMenB vaccine components was detected by ELISA in IP- and SC-immunized mice (Fig S1A), and IgG2a titers were higher in mice given 100 µL or 250 µL compared to 20 µL (Fig S1B). This dose response was mirrored in Western blots using anti-IgG secondary antibody against whole-cell lysates of *Nm* and six different strains *Ng* (Fig S1C). Serum from control mice given PBS or Alum adjuvant alone did not recognize any *Nm* or *Ng* proteins. No adverse effects were observed following IP injection. Nodules formed at the injection site in SC-immunized mice for all doses given, which resolved over time.

The MenNZB human epidemiology study reported by Petousis-Harris, et *al*. [26] was based on subjects who received a 3-dose regimen separated by one month. Upon demonstrating that the 250 µL dose of 4CMenB was well-tolerated and induced the highest serum antibody titers, we added a third immunization in subsequent mouse immunization/challenge experiments. Mice were given 250 μL of the formulated vaccine three times by IP or SC injection; controls received Alum or PBS (IP). A 3-week interval between immunizations was used to avoid increasing the age of the mice before challenge, which can reduce susceptibly to *Ng*. Significantly higher titers of *Ng*-specific serum total Ig, IgG1, and IgG2a (*p* < 0.004) but not IgA were detected in SC- and IP-immunized mice against *Ng* OMVs compared to control groups that received Alum or PBS on day 52 (Fig 1A-D). Total Ig, IgG1 and IgG2a titers were further elevated by the third immunization in both the IP- and SC-immunized groups (day 31 versus day 52) (p ≤ 0.05). The IgG1/IgG2a ratio was significantly lower for IP-immunized mice on day 31, but similar to SC-immunized mice on day 52 (Fig 1E) due to a marked increase in IgG2a titers in the SC-immunized group after the third immunization (Fig 1C). Vaginal total Ig and IgG1 (*p* < 0.0001), but not IgG2a or IgA were significantly elevated in vaginal washes collected after the second immunization compared to control groups in IP-immunized mice, but not SC-immunized mice (Fig 1F-I). Vaginal washes were not collected after the third immunization to avoid altering the vaginal microenvironment before bacterial challenge and cessation of the LeBoot effect, which would increase the number of mice in the undesired stages of the estrous cycle at the time of challenge [31]. We conclude that a half human-dose of 4CMenB is well-tolerated in mice and that a dosing regimen similar to that used in the New Zealand study elicits systemic and mucosal humoral immune responses that are cross-reactive against *Ng*.

**Fig 1.**
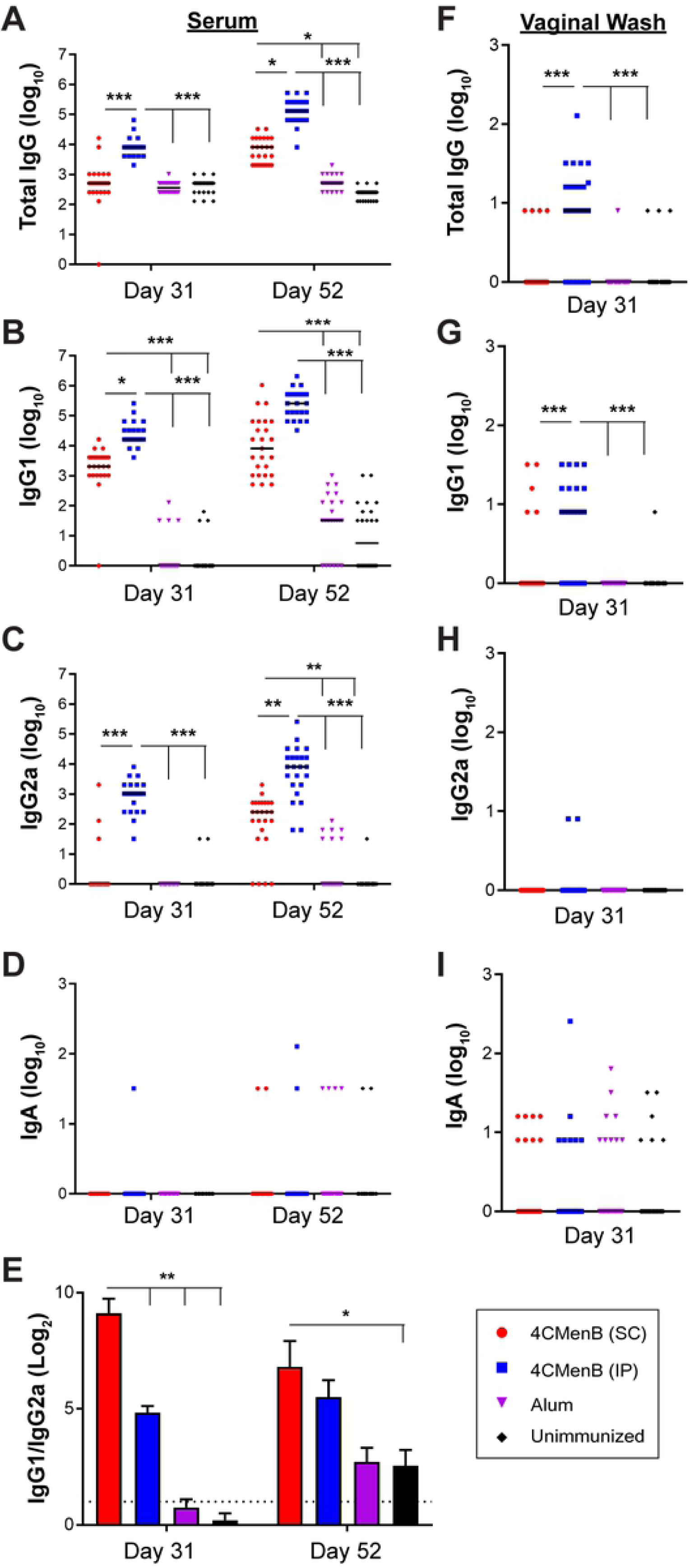
Serum and vaginal antibody titers against F62 OMVs from 4CMenB-immunized mice. Groups of 25 BALB/c mice were immunized three times, three weeks apart with 250 µl or 4CMenB by the IP or SC routes or with PBS or Alum (IP route). Serum and vaginal antibody titers on day 31 and day 52 (ten days after the 2^nd^ and 3^rd^ immunization, respectively) against F62 OMVs were measured by ELISA. Shown are serum **(A)** total Ig, **(B)** IgG1, **(C)** IgG2a, **(D)** IgA and **(E)** IgG1/IgG2a ratios for on days 31 and 52 (left column). Vaginal **(F)** total Ig, **(G)** IgG1, **(H)** IgG2a and **(I)** IgA on day 31 are shown in the right column. No difference was found in any sample or Ig tested between control animals receiving PBS and those receiving alum only. *, p<0.05; **, p<0.01; ***, p<0.0001. Results from the repeat experiment were similar.

### 4CMenB-immunized mice clear *Ng* infection significantly faster and have a reduced bioburden following vaginal challenge

To assess the protective efficacy of 4CMenB against *Ng*, we challenged 4CMenB-immunized and control mice with *Ng* strain F62 three weeks after the third immunization and quantitatively cultured vaginal swabs for *Ng* over seven days. In combined data from two independent experiments, IP-immunized mice exhibited a significantly faster clearance rate (p ≤0.0001) (Fig 2A) and lower bioburden compared to control groups given PBS or Alum alone (p <0.05 and ≤0.01, respectively) (Fig 2B and 2C). Data for each individual experiment, which also showed significantly faster clearance for both immunized groups compared to control mice, are shown in Fig S2A and S2C. The bioburden in the IP-immunized group was significantly lower than that of the Alum only group in both experiments, while the difference in the bioburden in SC-immunized mice compared to Alum was significant only in the repeat experiment (p < 0.0001) (Fig S2B and S2D). Combined data from the two experiments showed that 70% and 88% of mice given 4CMenB by the SC and IP routes, respectively, cleared infection by day 7 compared to 25-30% of mice given alum or PBS (Fig 2D).

**Fig 2.**
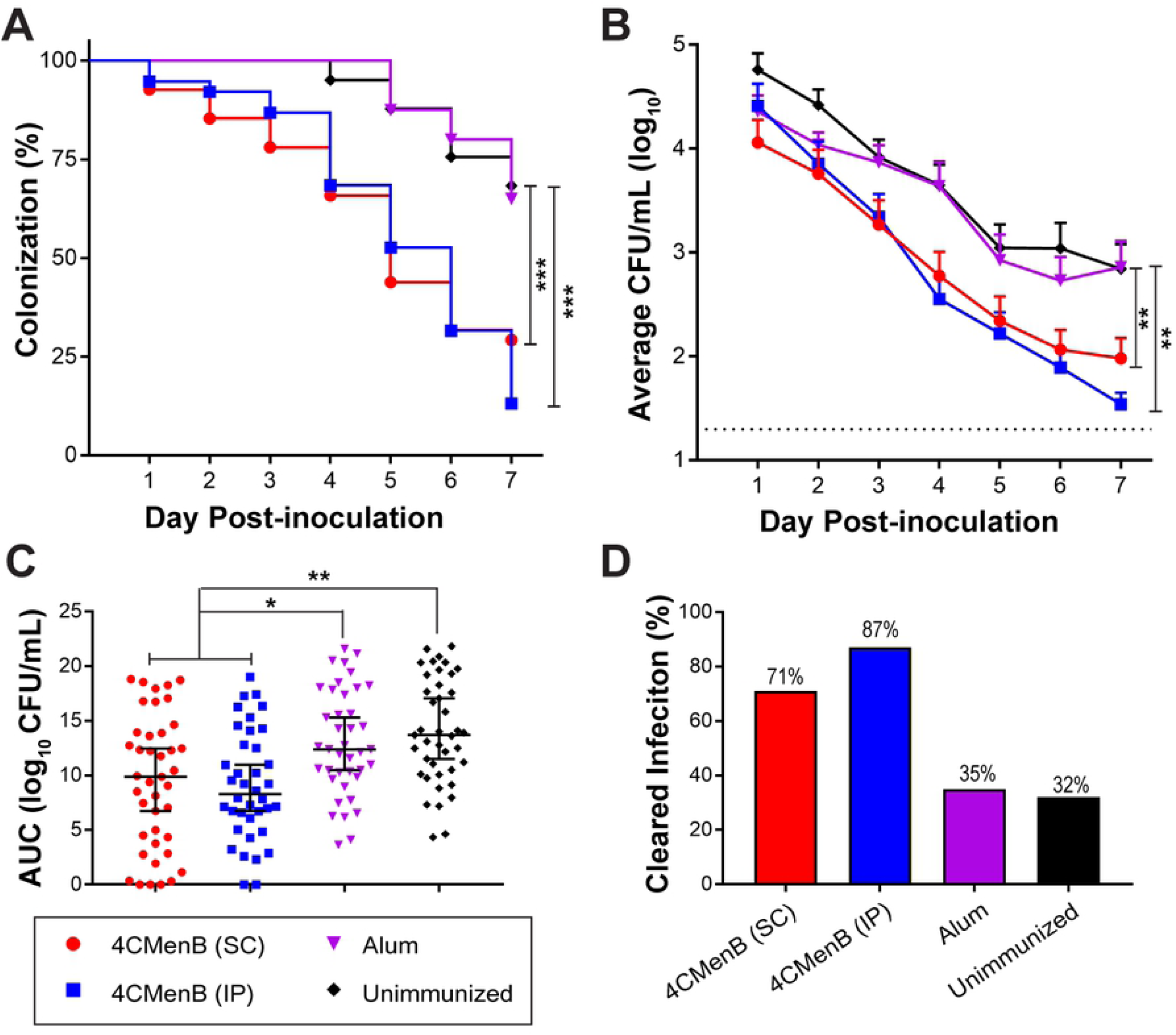
4CMenB has *in vivo* efficacy against *Ng*. Mice were immunized three weeks apart with 250-µL doses of 4CMenB by the IP (blue) or SC (red) route or with PBS (black) or alum (purple) by the IP route and challenged with *Ng* strain F62 three weeks after the final immunization. Shown are the combined data from two independent trials (total n = 38-41 mice/group). **(A)** Percentage of culture-positive mice over time; **(B)** Average CFU per ml of a single vaginal swab suspension; **(C)** total bioburden over 7 days expressed as area under the curve; **(D)** Percentage of mice that cleared infection by day 7 post-challenge. *, p < 05; **, p < 0.01; ***, p < 0.0001.

A peak vaginal PMN influx beginning on day 4 post-bacterial challenge was observed in all groups, and there was no difference in percentage of PMNs among experimental groups over time (Fig 3S). We also evaluated complement-dependent bactericidal activity of pooled sera from each group against *Ng* strain F62, the serum-sensitive challenge strain, and against the serum-resistant strain FA1090, using normal human serum as the complement source. The bactericidal_50_ titers were 1:480 and 1:240, respectively, which were 4-fold greater than that of pooled serum from the unimmunized group (Fig 3). We conclude that 4CMenB reproducibly accelerates clearance of *Ng* from the murine genital tract and lowers the bioburden over time and that opsonophagocytosis and complement-mediated bacteriolysis maybe contribute to the protection.

**Fig 3.**
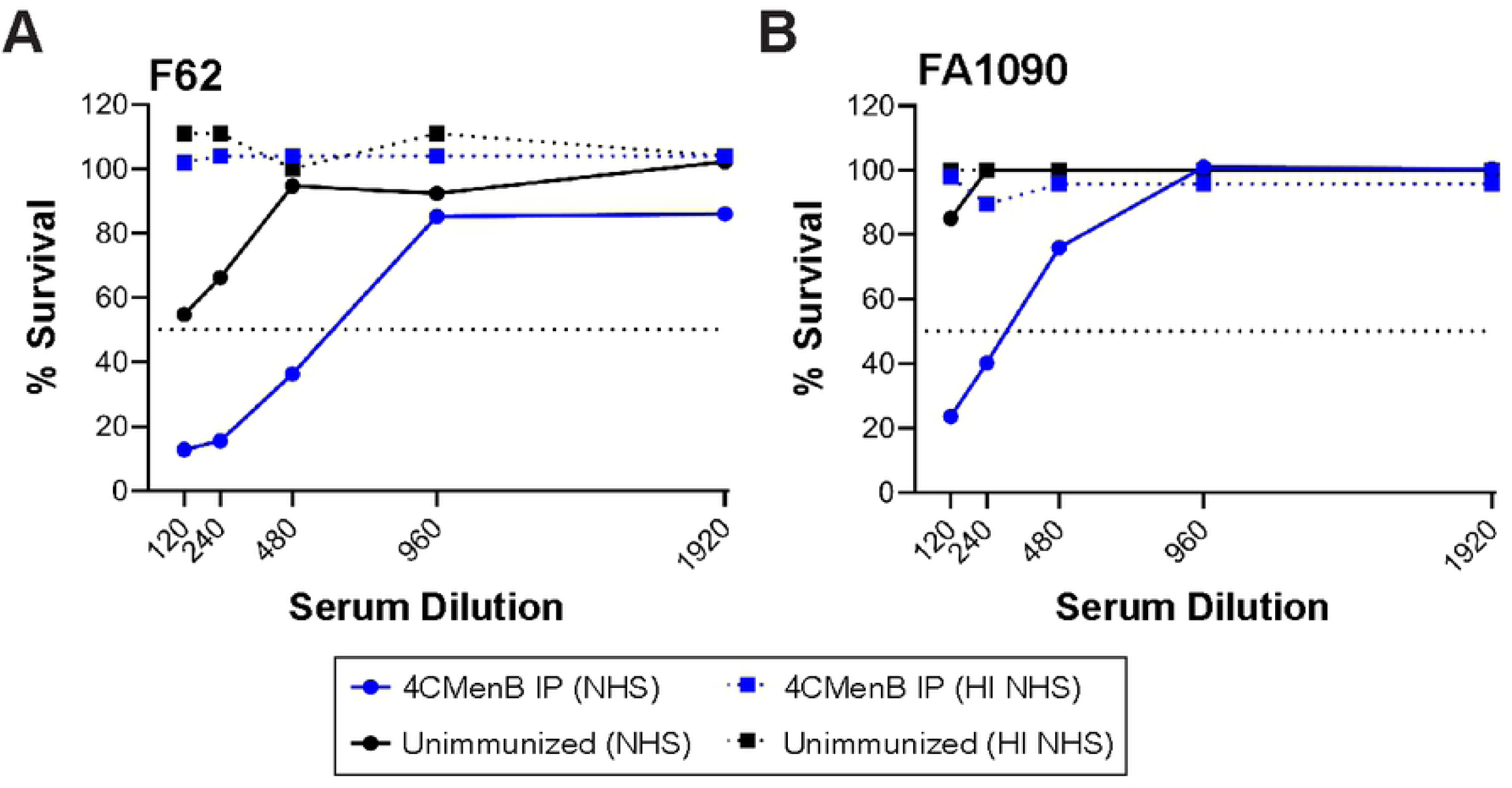
Antisera from 4CMenB-immunized mice is bactericidal against a serum-sensitive and a serum-resistant *Ng* strain. Serial dilutions of pooled serum from mice vaccinated with 250 µl of 4CMenB (blue lines) or Alum alone (black lines) by the IP route were incubated with 10^4^ CFU of the challenge strain F62 or strain FA1090 in microtiter plates as described in the Materials and Methods. After 5 min, NHS or heat-inactivated (HI)-NHS (final concentration, 10%) was added. After 45 min incubation at 37°C, the number of viable *Ng* in each well was determined by duplicate culture on GC agar. Data are expressed as the number of CFU from wells incubated with test serum divided by the number recovered from wells containing PBS instead of test serum, X times 100. Solid lines indicate NHS was used as the complement source; dotted lines represent data from wells tested in parallel with HI-NHS. The dotted line at 50% survival was drawn to identify the bactericidal_50_ titers.

### 4CMenB-Induced serum and vaginal antibodies cross-react with several *Ng* OMV proteins but not with *Ng* LOS species in a majority of strains

To examine the cross-reactivity of 4CMenB-induced antibodies against *Ng* surface proteins, we conducted western immunoblots against OMVs from the challenge strain and five other *Ng* strains that are geographically and temporally distinct in their isolation. Pooled antisera from mice immunized twice with the 250 μL dose by either the IP or SC routes (250IP or 250SC, respectively) recognized four prominent bands in fractionated OMV preparations all six strains: a high molecular weight (HMW) band > 220 kD, a doublet with bands of apparent molecular weight of 97 and 94 kDa, and a 55 kDa band (Fig 4). Several low intensity bands between 26 and 36 kDa were also recognized in several of the strains. Reactivity of the 250SC antiserum was weaker than the 250IP antiserum, which likely reflects the lower titers of this antiserum. Serum from PBS- or Alum-treated control mice did not recognize any *Ng* proteins (data not shown). Consistent with ELISA data, serum reactivity as assessed by band intensity was increased by a third immunization (Fig 4A), and additional bands were recognized including several bands in the 30-35 kDa range. A similar recognition pattern was observed on blots incubated with pooled vaginal washes and sera collected 10 days after the third immunization from immunized but unchallenged mice followed by anti-mouse IgG or anti-mouse IgA (Fig. 4B; compare lanes 1 and 2 with lane 5 in each blot). These results also show that while vaginal titers were low after the second immunization as measured by ELISA (Fig 1), vaccine-induced vaginal antibodies were readily detectable by immunoblot after a third immunization.

**Fig 4.**
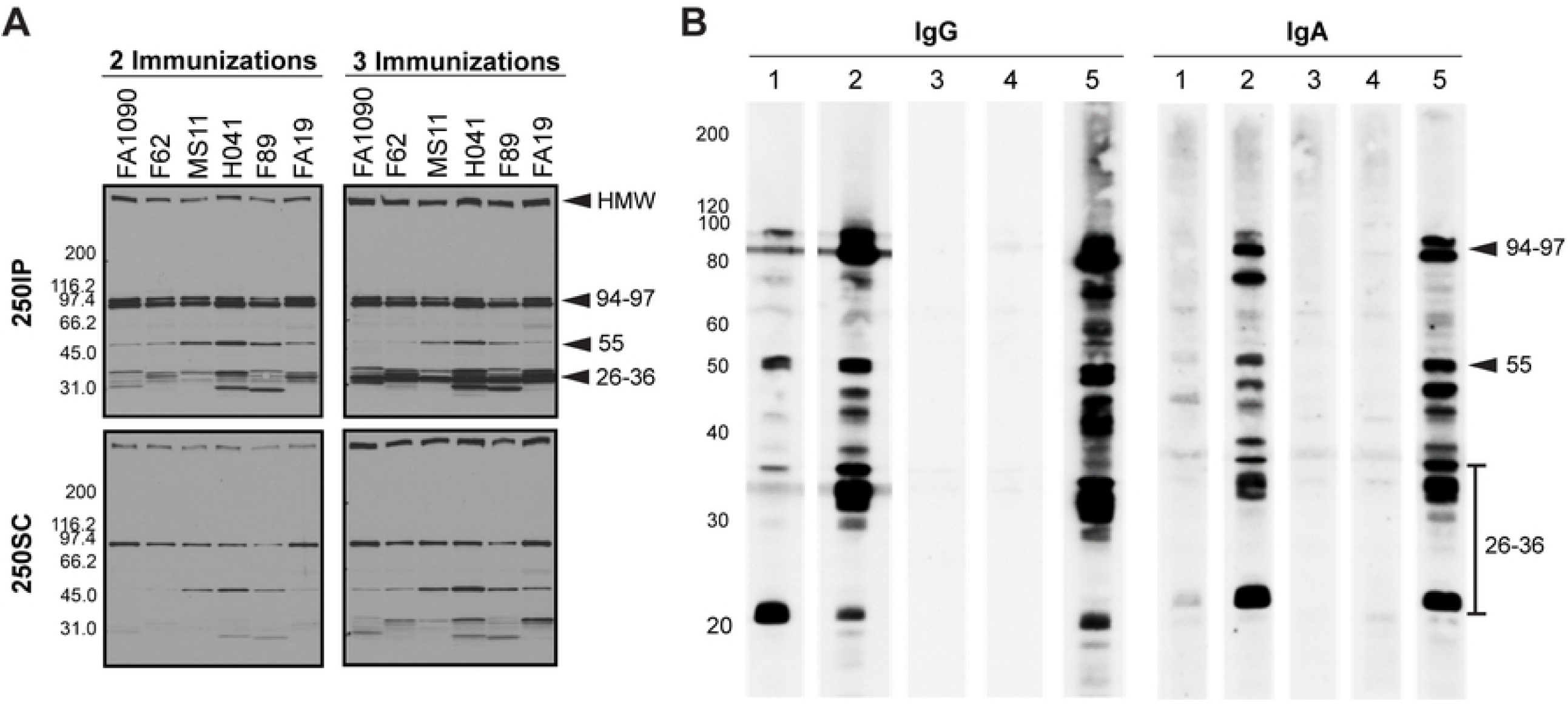
Serum and vaginal antibodies from 4CMenB-immunized mice recognize *Ng* outer membrane proteins by western immunoblot. **(A)** Pooled antisera from mice immunized with 250 µL of 4CMenB by the IP (250IP, upper panels) or SC (250SC, lower panels) route were tested against OMVs (app. 20 µg per lane) from 7 different *Ng* strains fractionated on 4-20% Tris-glycine gels by western blot (1:10,000 dilution of primary antisera) followed by secondary anti-mouse IgG-HRP. A boosting effect is observed when comparing the band intensities for serum collected after 2 and 3 immunizations. **(B)** Pooled vaginal washes from immunized or control mice collected after the third immunization tested against OMVs from the F62 challenge strain by western blot (1:100 dilution), followed by secondary anti-mouse IgG-HRP or anti-mouse IgA-HRP. Pooled vaginal washes were from mice given: (1) SC250; (2) IP250; (3) PBS; (4) Alum only. (5) Results using 250IP mouse serum (1: 10,000) for comparison. The band recognition pattern was similar for blots incubated with serum (lanes 5) or vaginal washes (lanes 1-4), and vaginal washes from IP-immunized mice were more strongly reactive than from SC-immunized mice. All lanes were equally loaded, as determined by Ponceau S staining. Shown are representative results from at least 2 separate experiments with identical results.

We also examined the reactivity of the immune serum against *Ng* lipooligosaccharide (LOS) based on the report that a significant percentage of the bactericidal activity induced by an *Nm* OMV vaccine was directed towards *Nm* LOS [32]. Neisserial LOS is a branched structure that consists of oligosaccharide extensions from the core oligosaccharide called the α- and β-chains.

An additional extension called the γ-chain is present in some strains [33]. Different LOS species can be produced within a strain due to phase variable expression of the glycosyltransferase genes *lgtA, lgtC, lgtD* and *lgtG*, which results in different lengths of the oligosaccharide chains [34, 35]. To test the cross-reactivity of 4CMenB antisera against *Ng* LOS, we fractionated crude LOS extracts by gel electrophoresis from four laboratory and thirteen clinical *Ng* isolates isolated between 1991 and 2019. Gels were stained or electroblotted to filters and incubated with monoclonal antibodies (3F11, 4C4 and 2C7) that recognize known epitopes within *Ng* LOS [36], or with the 4CMenB antiserum. All of the strains produced an LOS that bound one or more of monoclonal antibodies (Fig 5A). In contrast, the 4CMenB antiserum did not recognize the LOS in thirteen of the seventeen strains tested. The remaining four strains (MS11 and three clinical isolates LGB24, NMCSD322 and NMCSD6364) produced one or two LOS species that cross-reacted with the 4CMenB antiserum (Fig 5B). The lack of recognition of LOS in some strains could potentially be explained by the reactive LOS epitope being phase variable. Schneider *et al*. [37] demonstrated that long-chain LOS species are selected during urethral infection in men and Rice and colleagues have shown that the phase variable 2C7 LOS epitope is expressed among a majority of clinical isolates [36]. Therefore, to test investigate whether the anti-4CMenB-reactive LOS epitope is perhaps phase variable and selected *in vivo*, we infected mice with strain H041. No 4CMenB-reactive LOS species were detected in LOS preps from pooled vaginal H041 isolates cultured on days 2 and 5 post-inoculation (Fig 5C). We conclude that while cross-reactive antibodies to *Ng* LOS epitopes are induced by 4CMenB, the epitopes do not appear to be shared by a majority of *Ng* strains.

**Fig 5.**
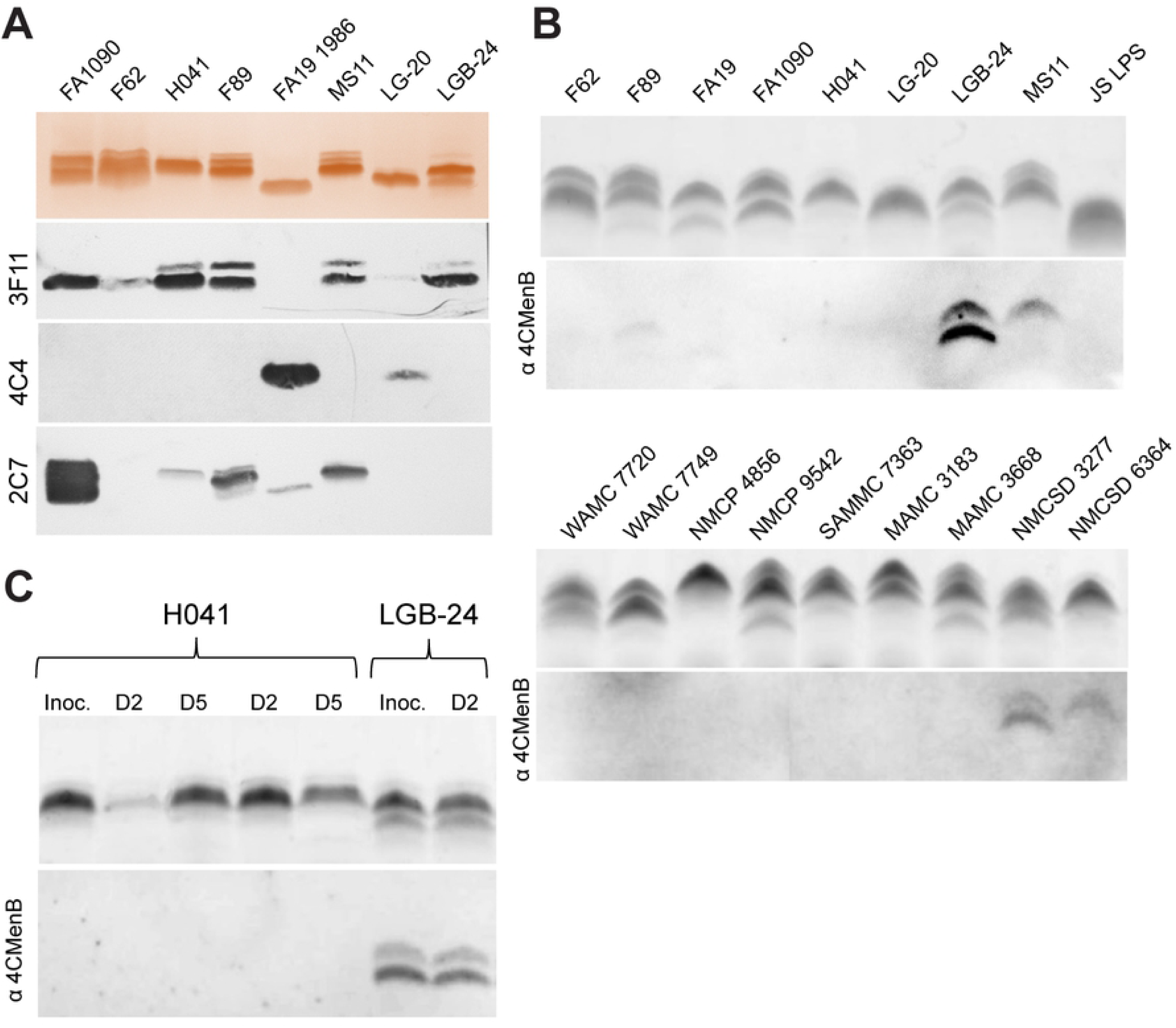
4CMenB antiserum recognizes *Ng* LOS in a minority of strains. Proteinase K-treated bacterial extracts from 4 laboratory strains and 13 clinical isolates were resolved on 16% Tricine gels and stained with (A) silver stain (top panel) or (B) Emerald green (top panels), or electroblotted and probed with the following: (A) Mabs 3F11, 4C4 and 2C7, which recognize *Ng* LOS epitopes (bottom panels). Note that FA19 1986 is a variant of FA19 (panel B) that has a phase-off *lgtA* gene that results in truncation of the LOS to a single 3.6 kDa species [91]. (B) Pooled IP250 4CMenB antisera (bottom panels). The doublets in LGB-24 and NMCSD 3277 and single LOS species in MS11 and NMCSD 6364 that were recognized by the antiserum are distinct from the LOS species identified by the Mabs shown in Panel A. (C) Emerald green-stained (upper panel) LOS from *Ng* strain H041 and LGB-24 (positive control) used to inoculate mice (Inoc) and from vaginal cultures collected on days 2 and 5 of infection. No change in the LOS species or 4CMenB reactivity was observed during infection by these strains.

### 4CMenB induces antibodies against promising *Ng* vaccine targets

To identify the proteins recognized by 4CMenB-induced antisera, we fractionated OMVs from *Ng* strain F62 on two separate gels. One was stained with a G-250 Coomassie stain for mass spectrometry analysis, while the other was used for Western blotting with the 250IP antiserum. The blot and gel were aligned and the reactive bands identified by molecular weight and band intensity. Bands indicated by the numbered arrows (Fig 6A), which correspond to the most intensely recognized bands in the Western blot (Fig. 6B), were submitted for mass spectrometry analysis (Table 1). The HMW band at the top of the gel was identified as PilQ, which is a protein that forms a dodecamer through which gonococcal pili extend [38]. Mass spectrometry analysis identified two potential proteins in band 2 (97 kDa): an elongation factor and a phosphoenolpyruvate. Band 3 (94 kDa) also contained 2 potential proteins: BamA, an Omp85 homologue involved in the biogenesis of OM proteins (OMPs) [39], and a methyltransferase. Band 4 (55 kDa) was identified as MtrE, the OM channel of the three different gonococcal active efflux pump systems [40]. The 36 kDa protein (band 5) was identified as PorB, and the 32 kDa band (band 6), as Opa. In summary, we identified eight proteins from six cross-reactive bands, five of which are known surface-exposed *Ng* antigens.

**Table 1.**
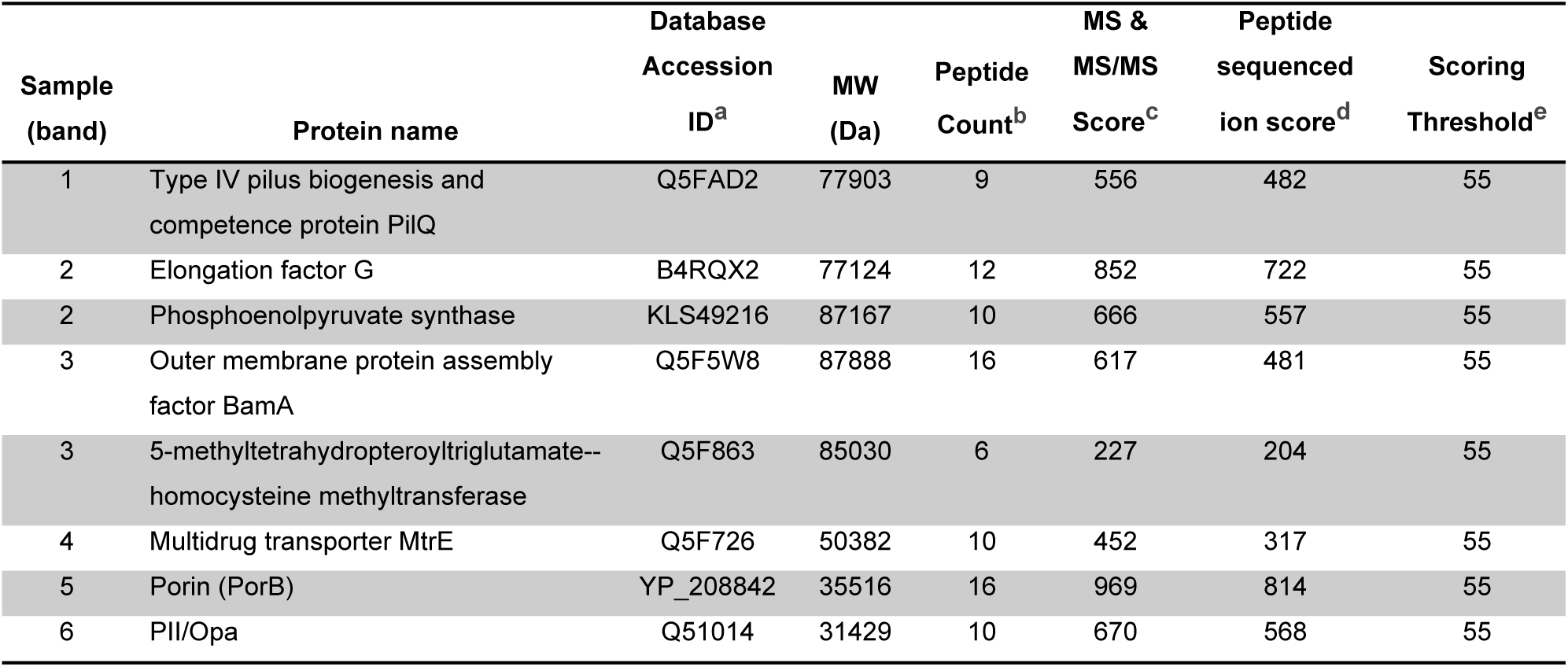
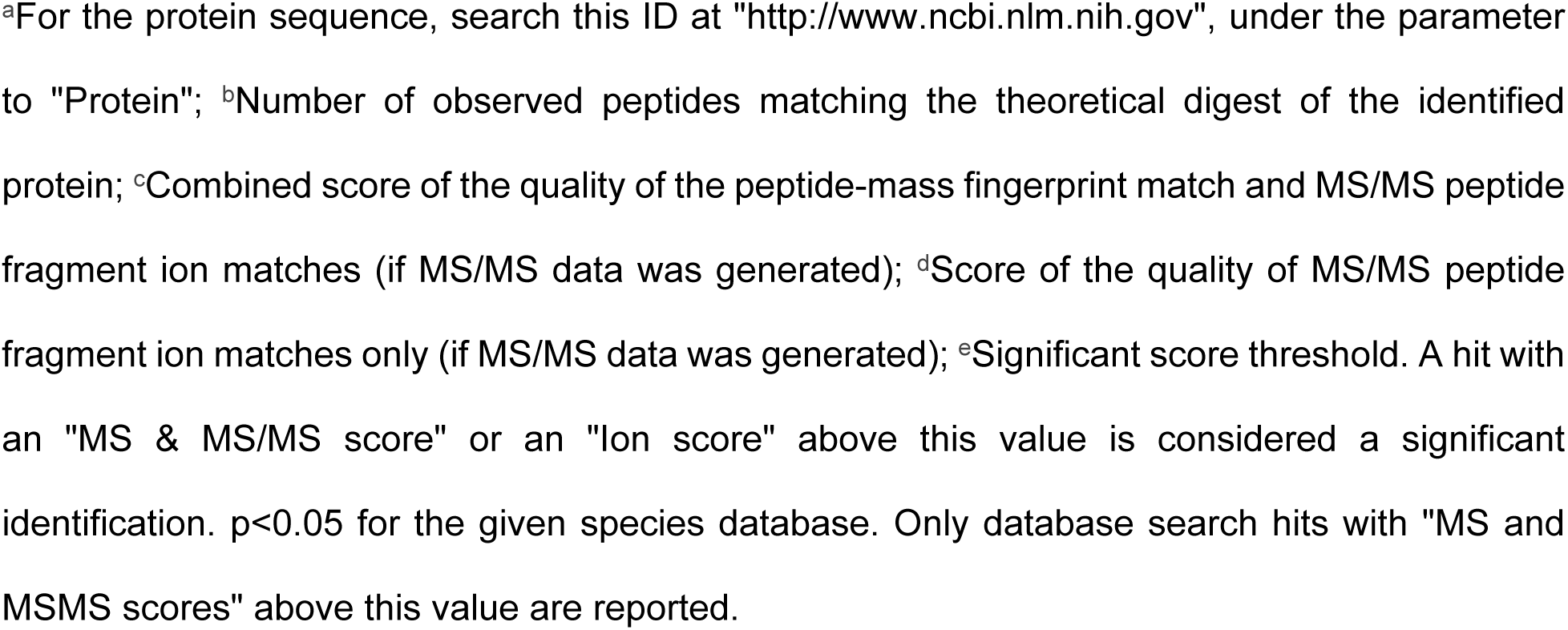
Identification of protein bands recognized by 4CMenB antisera as determined by mass spectrometry.

**Fig 6.**
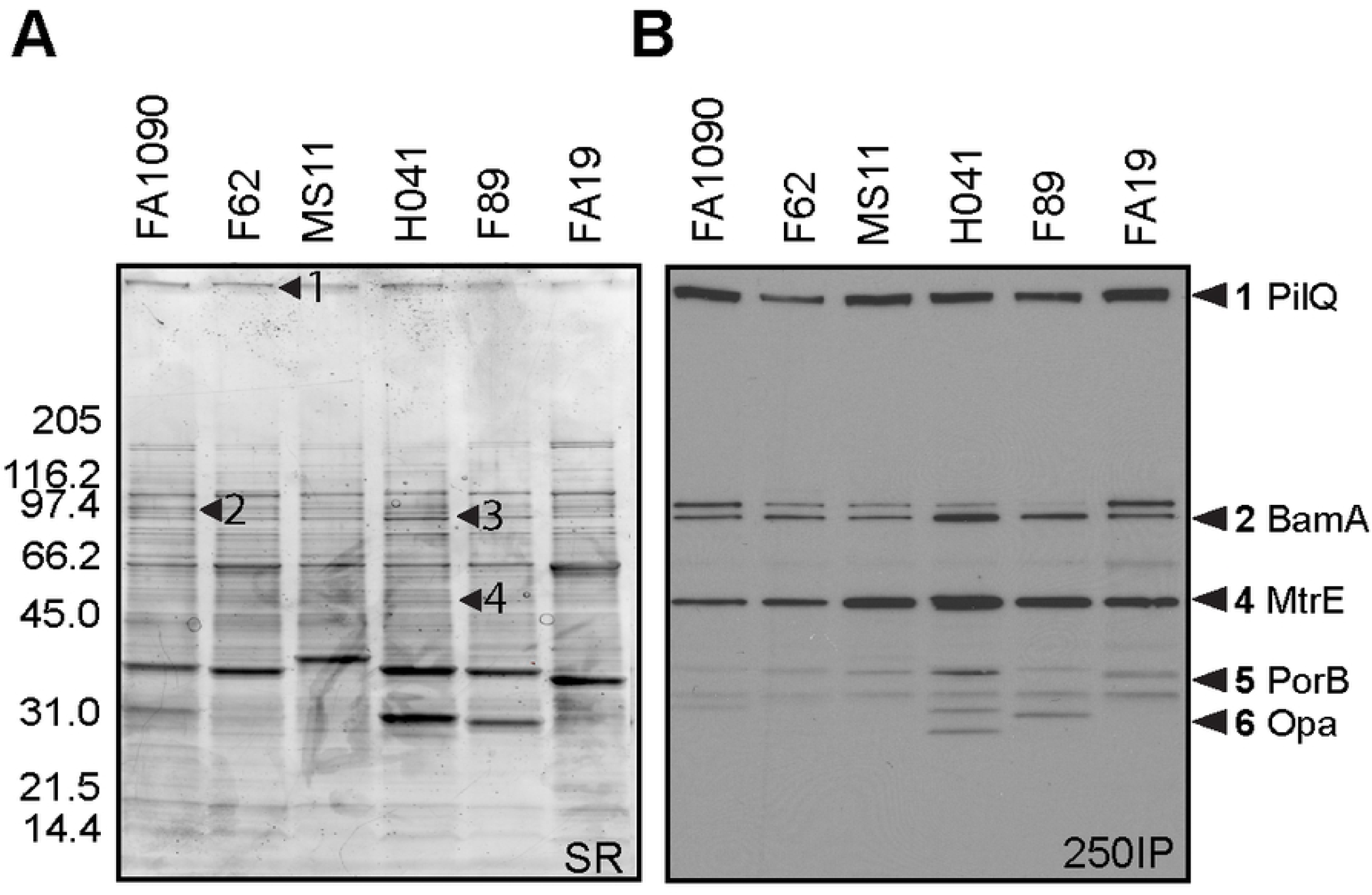
PilQ, MtrE, porin and OpA are recognized by 4CMenB antisera. OMV (app. 20 µg) from 6 *Ng* strains (Table 2) were subjected to SDS-PAGE on a 4-20% Tris-glycine gel and (A) stained with sypro ruby or (B) transferred to PVDF for western blot with the 250IP antiserum. The stained gel was aligned with the Western blot, and corresponding bands were digested and analyzed by mass spectrometry. The numbers indicated with arrows on each panel correspond to the same numbers on the Western blots except for bands 5 and 6, which were excised from a different gel but are indicated on the western based on the banding patterns. Proteins identified are described in Table 1. Among the proteins identified, known surface-exposed outer membrane proteins are: (1) PilQ, (2) BamA, (4) MtrE, (5) PorB, and (6) Opa.

### 4CMenB-induced antibodies bind native PilQ and MtrE on the surface of viable gonococci

We next performed immunoprecipitations using 4CMenB-induced mouse antisera and live FA1090 and MS11 bacteria. Antigen-antibody complexes were solubilized in detergent, retrieved using protein A/G agarose, and subjected to non-denaturing Western blotting with the 250IP antiserum. Whole cell lysates and OMVs were run in parallel and exhibited the same reactive band pattern shown in Figs 4A and 6B. Serum from the alum-alone group pulled down non-specific proteins smaller than 30 kDa that did not align with bands in lanes containing whole cell lysates or OMVs. In contrast, the 250IP antiserum pulled down two proteins from both strains: a HMW protein, possibly PilQ, and a ∼ 55 kDa protein (Fig 7). We hypothesized the 55 kD protein was MtrE based on the greater intensity of this band in western blots against OMVs from strains MS11, H041 and F89, which carry one or more *mtr* mutations that cause increased production of the MtrCDE efflux pump (Fig 4A).

**Table 2.**
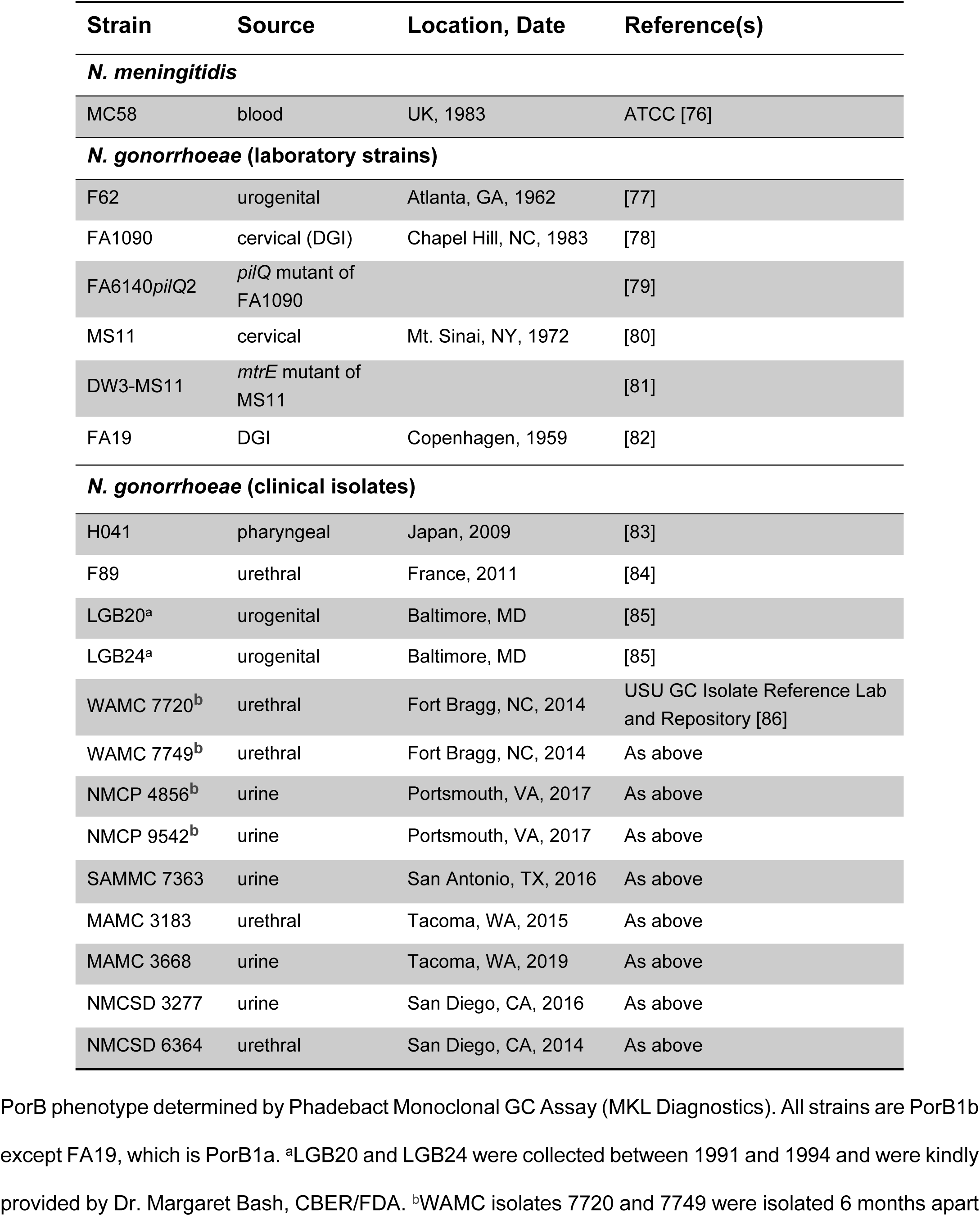
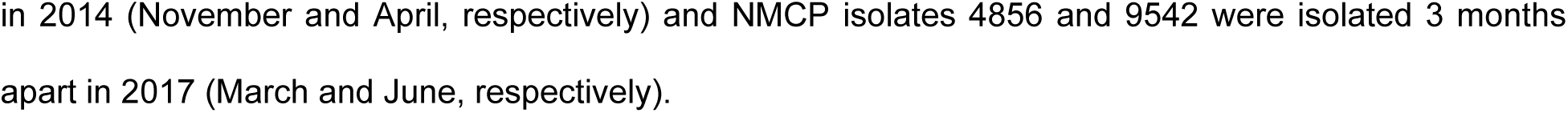
Bacterial strains used in this study.

| Strain | Source | Location, Date | Reference(s) |
| --- | --- | --- | --- |
| ***N. meningitidis*** |  |  |  |
| MC58 | blood | UK, 1983 | ATCC [76] |
| ***N. gonorrhoeae* (laboratory strains)** |  |  |  |
| F62 | urogenital | Atlanta, GA, 1962 | [77] |
| FA1090 | cervical (DGI) | Chapel Hill, NC, 1983 | [78] |
| FA6140*pilQ*2 | *pilQ* mutant of FA1090 |  | [79] |
| MS11 | cervical | Mt. Sinai, NY, 1972 | [80] |
| DW3-MS11 | *mtrE* mutant of MS11 |  | [81] |
| FA19 | DGI | Copenhagen, 1959 | [82] |
| ***N. gonorrhoeae* (clinical isolates)** |  |  |  |
| H041 | pharyngeal | Japan, 2009 | [83] |
| F89 | urethral | France, 2011 | [84] |
| LGB20<sup>a</sup> | urogenital | Baltimore, MD | [85] |
| LGB24<sup>a</sup> | urogenital | Baltimore, MD | [85] |
| WAMC 7720<sup>b</sup> | urethral | Fort Bragg, NC, 2014 | USU GC Isolate Reference Lab and Repository [86] |
| WAMC 7749<sup>b</sup> | urethral | Fort Bragg, NC, 2014 | As above |
| NMCP 4856<sup>b</sup> | urine | Portsmouth, VA, 2017 | As above |
| NMCP 9542<sup>b</sup> | urine | Portsmouth, VA, 2017 | As above |
| SAMMC 7363 | urine | San Antonio, TX, 2016 | As above |
| MAMC 3183 | urethral | Tacoma, WA, 2015 | As above |
| MAMC 3668 | urine | Tacoma, WA, 2019 | As above |
| NMCSD 3277 | urine | San Diego, CA, 2016 | As above |
| NMCSD 6364 | urethral | San Diego, CA, 2014 | As above |

PorB phenotype determined by Phadebact Monoclonal GC Assay (MKL Diagnostics). All strains are PorB1b except FA19, which is PorB1a. <sup>a</sup>LGB20 and LGB24 were collected between 1991 and 1994 and were kindly provided by Dr. Margaret Bash, CBER/FDA. <sup>b</sup>WAMC isolates 7720 and 7749 were isolated 6 months apart
in 2014 (November and April, respectively) and NMCP isolates 4856 and 9542 were isolated 3 months apart in 2017 (March and June, respectively).

**Fig 7.**
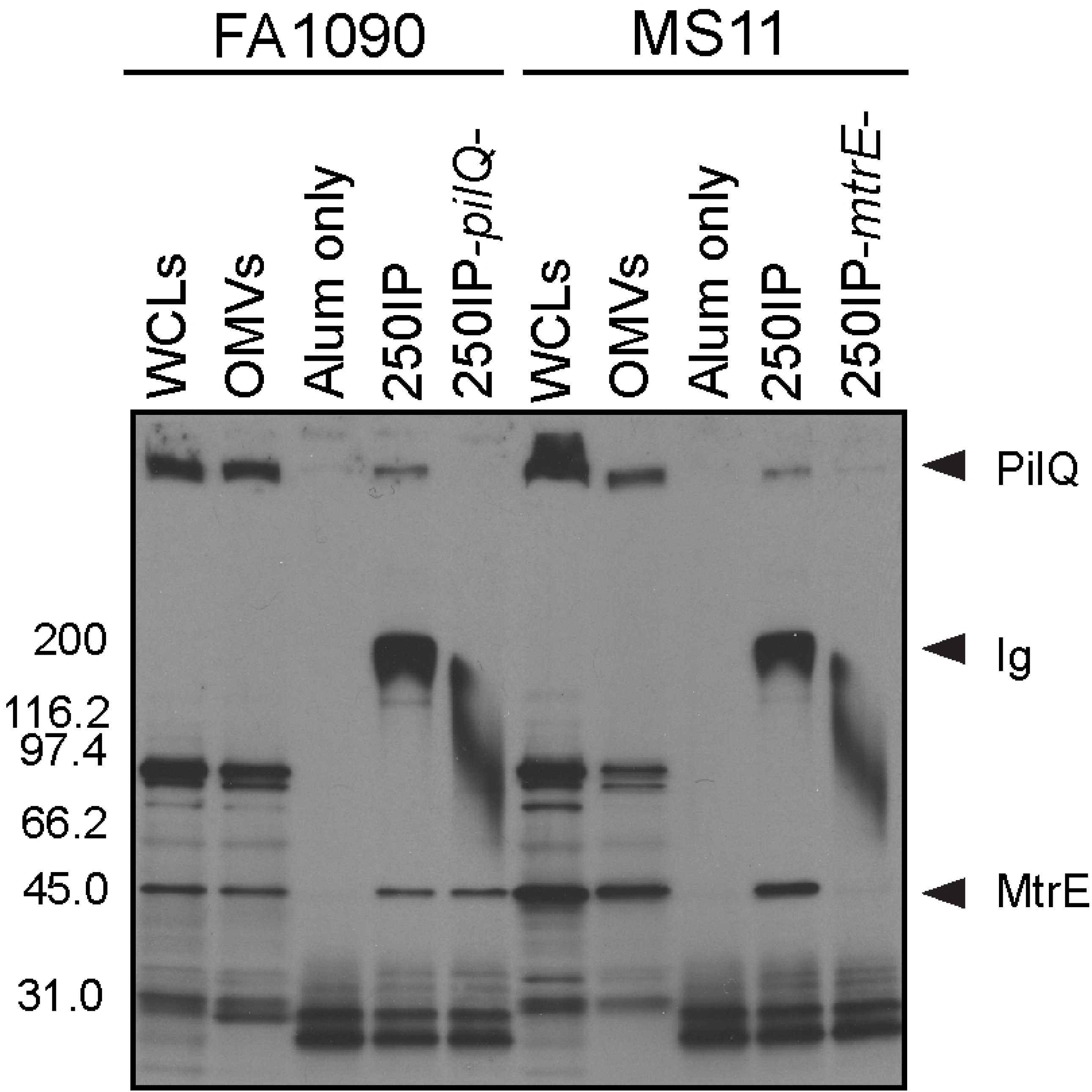
4CMenB -induced antibodies bind PilQ and MtrE at the surface of viable *Ng* FA1090 and MS11 bacteria. Immunoprecipitations were performed with wild-type strains FA1090 and MS11 and their isogenic *pilQ* and *mtrE* mutants using antisera from mice immunized with 250 µL of 4CMenB via the IP route (250IP) or given alum only (negative control). Bacterial components bound by 4CMenB-induced antisera were subjected to SDS-PAGE (non-denaturing conditions, 4-20% Tris-glycine) and Western blotting with 4CMenB 250IP antiserum. WCLs, total cellular proteins; OMVs, outer membrane vesicles; *pilQ*-, FA1090Δ*pilQ*; *mtrE*-, MS11Δ*mtrE*. Data shown are representative of at least 2 separate experiments with identical results. The wide band around 200 kDa corresponds to the antibodies within the test antisera that are present in the antigen-antibody complexes and pulled down with the protein A/G agarose.

To confirm that the two identified proteins are PilQ and MtrE, we included single isogenic mutants that lack expression of either protein, and the results confirmed the identity of the two immunoprecipitated proteins as PilQ and MtrE. The HMW band but not the 55-kDa protein was absent from the sample in which mutant strain FA1090*pilQ* was incubated with the antisera. Inversely, immunoprecipitation with mutant MS11*mtrE-* did not yield a band around 55 kDa, but did retain binding to the HMW protein PilQ. We conclude that 4CMenB induces antibodies recognize the native conformation of *Ng* PilQ and MtrE on the gonococcal surface.

## Discussion

The pathogenic *Neisseria* are human-specific pathogens that differ in the capacity to cause life-threatening septicemia and meningitis (*Nm*) and nonulcerative sexually transmitted infections of the urogenital tract that can ascend to cause damage to the upper reproductive tract (*Ng*). The only reservoir for these pathogens is human pharyngeal, genital and rectal mucosae where the bacteria reside extracellularly and within an intracellular niche [41, 42]. *Nm*, unlike *Ng*, produces a complex polysaccharide capsule that is critical for invasive disease, and vaccines that target the capsules of four of the five most prevalent capsular serogroups A, C, W135 and Y have been effectively used for decades. This approach is not successful against serogroup B *Nm* due to the α2-8-linked polysialic acid composition of the serogroup B capsule, which mimics α2-8-sialylated human glycoproteins [43]. A recent advance in biomedical research was the development of two licensed vaccines that prevent serogroup B *Nm* invasive disease, one consisting of purified lipoprotein subunits, rLP2086 (Trumenba®, Pfizer), and the other, 4CMenB (Bexsero®, GSK) [43]. The 4CMenB vaccine was proceeded by *Nm* OMV vaccines that were tailor-made against endemic serogroup B strains in New Zealand, Brazil, Cuba and Norway [27, 43].

Like vaccine development for serogroup B *Nm*, gonorrhea vaccine research has focused on conserved outer membrane proteins, *Ng* outer membrane vesicles, and in one case, the 2C7 oligosaccharide epitope within *Ng* LOS [18]. Vaccine development for gonorrhea is more complicated, however, by a lack of defined immune correlates of protection. Meningococcal vaccine development is guided by serum bactericidal activity, which was first demonstrated as the main correlate of protection against *Nm* invasive disease in a classic case-control study at Fort Ord, California in the 1960s [44]. Natural history studies for gonorrhea, in contrast, have led only to an association between antibodies against the restriction modifiable protein (Rmp) and increased susceptibility to infection, and in high-risk women, antibodies to porin and Opa proteins as being associated with reduced risk of *Ng* upper reproductive tract infection [18]. The lack of clear correlates of protection and the absence of immunity to reinfection have challenged the possibility of a gonorrhea vaccine.

It is in this context that the reported reduced risk of gonorrhea in subjects immunized with an *Nm* OMV vaccine [26] may herald a breakthrough for gonorrhea vaccine development. In support of this epidemiological evidence, we demonstrated that 4CMenB reproducibly accelerated *Ng* clearance and lowered the bioburden of *Ng* in a well-characterized mouse model of genital tract infection. 4CMenB induced vaginal and serum IgG1, IgG2a and IgA when administered subcutaneously that cross-react with several *Ng* OMV proteins expressed by six different *Ng* strains. The dosing regimen we used was similar to that of the MenNZB vaccine given to the subjects in the retrospective case-control epidemiological study [26]. These data are direct evidence of cross-species protection and suggest female mice may reproduce vaccine-induced mechanisms that protect humans against gonorrhea.

Known host-restrictions that limit the capacity of mice to mimic human neisserial infections have been extensively reviewed [18]. Restricted factors include receptors for several neisserial colonization or invasion ligands that mediate adherence to and, or uptake by human cells, including the type IV pili, the Opa proteins, and in *Ng*, the pili/PorB/C3b complex. In experimental murine infection, *Ng* is seen adherent to vaginal epithelial cells, in cervical tissue and within the lamina propria, which is presumably mediated by other *Ng* colonization ligands for which species specificity has not been defined [45, 46] or that are not host-restricted [47]. Acquisition of iron from lactoferrin (LF) and transferrin (TF), and recently, zinc from calprotectin [48], is also host-restricted. These restrictions limit the ability to fully evaluate the efficacy of vaccines that induce antibodies that block colonization or nutrient uptake in the mouse model, although mice that are transgenic for the human carcinoemybryonic antigen cellular adherence molecules (CEACAMs), the major Opa protein receptors and hTF could be used [49-51]. Restrictions in soluble negative regulators of the complement cascade, factor H (fH) and C4b-binding protein (C4BP), also exist and are especially important to consider when testing vaccines that clear *Ng* infection through bactericidal and opsonophagocytic activity. Transgenic hFH and hC4BP mice made for this purpose and not used in our study, were recently utilized by Rice and colleagues to more rigorously test the *in vivo* efficacy of immunotherapeutic strategies against *Ng* [52].

While unable to fully mimic human neisserial infections, animal models provide a physiologically relevant and immunologically intact system for testing vaccine-induced immune responses against infection. Meningococcal vaccine development is aided by the use of mice or rabbits to test whether candidate vaccines induce bactericidal antibodies against *Nm* or cause adverse effects [53, 54], and improved mouse and infant rat bacteremia models have been used to measure the efficacy of candidate *Nm* vaccines in eliminating *Nm* from the bloodstream [50, 55, 56]. Early gonorrhea vaccine studies used chimpanzees, which do not have all the host restrictions found in other animal species, and human male subjects [18]. Chimpanzees are no longer used for gonorrhea research, however a human urethritis model is still available [57] and is the most relevant model for studying vaccine efficacy against *Ng* urethral infection.

Currently, the estradiol-treated mouse model is the only animal model for studying gonorrhea vaccine efficacy in females, where the majority of morbidity and mortality associated with gonorrhea occurs. This model is also used to systematically screen antigens, immunization regimens, and adjuvants, and to analyze host responses [18]. Similarities between human and experimental murine infection include the fact that mice, like humans, produce a transient and unremarkable humoral response to *Ng* infection and can be reinfected with the same strain. *Ng* induces the Th17 pathway in both humans [19, 20] and mice [58, 59], which leads to recruitment of PMNs to the infection site. Similar to that reported for human cervical infections, hormonally driven, cyclical fluctuations in *Ng* colonization load and selection for Opa protein phase variants occurs over the course of murine infection [60]. *Ng* is seen within murine PMNs and importantly, *Ng* mutants that are more or less susceptible to killing by human PMNs or cationic antimicrobial peptides *in vitro* have a similar phenotype when tested against murine PMNs and cathelicidins, and are more fit or attenuated compared to the wild-type strain, respectively, during murine infection [60, 61].

We also showed that the 4CMenB vaccine induced serum bactericidal activity and high titers of serum IgG1 and IgG2a that cross-reacted with *Ng* OMVs. Vaccine-induced vaginal IgA was readily detected by immunoblot. Recruitment of PMNs to the infection site occurred in all groups, which could enable opsonophagocytic killing of *Ng* in the presence of specific antibody. Serum bactericidal activity was also detected that could mediate protection through complement-mediated bacteriolysis. Detailed investigation of the mechanism of protection have thus far only been reported for one candidate *Ng* OMV vaccine [62] and the 2C7 vaccine [36], both of which induced Th1 responses and bactericidal antibodies. The demonstration by Russell and colleagues that a vaginally applied Th1-inducing cytokine adjuvant clears *Ng* infection in mice and induces a specific adaptive response and memory humoral response also suggests Th1 responses are protective [63]. Whether this is true for human infection is not known. The importance of bactericidal activity in clearing *Ng* mucosal infections, is also not known and may be vaccine-specific. Passive protection studies with bactericidal monoclonal antibody against the 2C7 epitope clearly showed antibodies were sufficient for vaccine-mediated clearance [36]; however, eleven other promising purified protein subunit vaccines that induced high-titered bactericidal activity against *Ng* have been tested by our laboratory over the years that did not show protective efficacy in the gonorrhea mouse model (A.E. Jerse, unpublished data in collaboration with others). Further investigation of the immune responses induced by 4CMenB in mice and vaccinated humans should define mechanisms of vaccine-mediated protection. The use of 4CMenB as a commercially available positive control will also strengthen the protocol for screening of candidate gonorrhea vaccine antigens in the mouse model.

Recently, the proteome of the 4CMenB vaccine was defined and shown to contain 461 proteins, of which 60 proteins were predicted to be inner membrane or periplasmic and 36 were predicted to be in the outer membrane or extracellular [64]. Others identified twenty-two *Nm* proteins as comprising >90% of the 4CMenB proteome, twenty of which have homologs in *Ng*, These investigators also showed that post-vaccinated serum from 4CMenB-immunized humans recognized several bands in fractionated *Ng* OMVs by immunoblot [30]. In our study, the 4CMenB-induced antisera recognized several denatured OMV proteins in a panel of diverse *Ng* strains, which is consistent with the OMV portion of the 4CMenB vaccine generating a robust cross-reactive response. Using mass spectrometry, we identified eight cross-reactive proteins, including MtrE, PilQ and BamA, which are promising vaccine candidates [18]. Recognition of LOS species was less consistent, with only ca 25% of strains expressing LOS species that were reactive with anti-4CMenB serum. We also demonstrated that 4CMenB-induced antibodies bound native PilQ and MtrE at the surface of viable *Ng*.

The 94% amino acid identity (S1 Table) between the *Nm* and *Ng* homologs identified in this report is consistent with the cross-reactivity that we observed. Importantly, the residues encompassing the two short surface-exposed loops of the MtrE monomer (residues 92-99 and 299-311 [65]), which are highly conserved among *Ng* strains, are identical to those from MtrE expressed by *Nm* strain MC58 (data not shown). MtrE is the outer membrane channel of the MtrCDE, FarABMtrE and MacABMtrE efflux pumps, which expel antibiotics and host-derived antimicrobial compounds [66]. The importance of the MtrCDE active efflux pump in protecting *Ng* against host innate effectors has been demonstrated in the mouse model [67]. Antisera directed against the two surface-exposed MtrE loops could target *Ng* for complement-mediated bacteriolysis and opsonophagocytosis, and may possibly impair efflux pump function to increase *Ng* susceptibility to host innate effectors. The PilQ protein is critical for pilus secretion [68] and mutations in PilQ are associated with increased entry of heme and antimicrobial compounds [69] and enhanced resistance to cephalosporin [70]. Amino acids 406 to 770 of *Nm* PilQ were shown to be a promising vaccine target, and are 94% identical with the same region of *Ng* PilQ (strain FA1090) [71]. BamA is a surface-exposed outer membrane belonging to the Omp85 family [72, 73]. The essential role of BamA in outer membrane protein biogenesis suggests it may be a highly effective vaccine target as it is present in cell envelopes and OMVs, surface-exposed, and is well-conserved among clinical *Ng* isolates [39].

In summary, the demonstration that a licensed *Nm* OMV-based vaccine accelerates *Ng* in clearance in a genital tract infection mouse model is direct evidence that cross neisserial species protection may be an effective vaccine strategy for gonorrhea. Whether this approach would protect against *Ng* rectal or pharyngeal infections, which are very common, is not known and in the absence of animal or human challenge models for these infections, this question must be solely addressed by epidemiological or clinical trials. Future detailed immunological studies in mice, which can be experimentally manipulated to directly test hypothesized mechanisms of protection, combined with clinical research studies on 4CMenB-vaccinated humans should reveal new and important information on how to combat this ancient, highly successful pathogen.

## Materials and Methods

### Bacterial strains and culture conditions

*Ng* strains used in this study are listed in Table 2. Supplemented GC agar (Difco) was used to routinely propagate *Ng* as described [74]. GC-VNCTS agar [GC agar with vancomycin, colistin, nystatin, trimethoprim (VCNTS supplement; Difco) and 100 µg/ml streptomycin (Sm)] and heart infusion agar (HIA) were used to isolate *Ng* and facultatively anaerobic commensal flora, respectively, from murine vaginal swabs [75].

### Immunizations and challenge experiments

Four-week-old female BALB/c mice (Charles River; NCI Frederick strain of inbred BALB/cAnNCr mice, strain code 555) were used in these studies. In pilot immunization studies, groups of 5 mice each were immunized with 20, 100 or 250 µL of 4CMenB (GSK) by the intraperitoneal (IP) or subcutaneous (SQ) routes on days 0 and 28. Two independent immunization and challenge experiments were conducted. For these experiments, 250 µL of the vaccine were given IP or SC on days 0, 21 and 42. Control mice received PBS or alum in the form of Alhydrogel (InVivogen) diluted in PBS (n = 20-25 mice/group). Venous blood was collected on days 31 and 52; vaginal washes were collected on day 31. Three weeks after the final immunization, mice in the anestrus or the diestrus stage of the reproductive cycle were implanted subcutaneously with a 21-day slow-release 17β-estradiol pellet (Innovative Research of America) and treated with antibiotics to suppress overgrowth of potentially inhibitory flora as described [60]. Two days after pellet implantation, mice were inoculated vaginally with 10^6^ colony-forming units (CFU) of *Ng* strain F62. Vaginal swabs were quantitatively cultured for *Ng* on 7 consecutive days post-challenge and used to prepare stained smears to examine the influx of vaginal polymorphonuclear leukocytes (PMNs) [75].

### Enzyme-linked immunosorbent assay (ELISA) and western blots

Serum or vaginal total Ig, IgG1, IgG2a and IgA were measured as endpoint titers as determined by standard ELISA [87]. Microtiter plates were coated with 20 µL/well of a 1:5 dilution of the formulated Bexsero vaccine in 15 mM Na_2_CO_3_, 35 mM NaHCO_3_, pH 9.5, or with 4 µg/ml of OMV from *Ng* strain F62. OMVs were isolated from supernatants from late-logarithmic phase cultures that were centrifuged for 1 hour at 100,000 x g at 4 °C. Pellets were resuspended in 1 mL of PBS. Protein concentration was determined by the BCA protein assay (Thermo Scientific). For whole-cell lysates (WCL) (total cellular proteins), bacteria from agar plates or mid-logarithmic phase cultures were centrifuged and the bacterial pellets suspended to an OD_600_ = 0.5. One milliliter of this suspension was mixed with 60 µL of Laemmli sample buffer. For western blots, WCL (4 µL) or 20 µg of OMV were subjected to sodium dodecyl sulfate polyacrylamide gel electrophoresis (SDS-PAGE), transferred to nitrocellulose membranes, stained with Ponceau S, and blocked overnight with 0.5% Tween20 in PBS. Membranes were incubated with pooled antisera or vaginal washes from each experimental group diluted in block and washed three times with 0.05% Tween 20 in PBS. Secondary antibody was horseradish peroxidase (HRP)-conjugated anti-mouse IgG or IgA and a chemiluminescence HRP was used as substrate (GE Healthcare). Apparent molecular weight of bands was determined using a standard curve generated from molecular weight markers (r^2^=0.9916; http://www.bio-rad.com/webroot/web/pdf/lsr/literature/Bulletin_6210.pdf). For western blots with LOS, proteinase K-treated bacterial extracts were generated as described [88] without the phenol treatment step, separated on 16% tricine gels (Novex) and probed with 1:10,000 dilutions of pooled serum from immunized and control mice followed by HRP-conjugated anti-mouse IgG as above.

### Immunoprecipitation and mass spectrometry

Two milliliters of a *Ng* suspension (OD_600_ = 1) prepared from a mid-logarithmic phase Gc broth culture were mixed with 30 µL of antisera for 20 minute at room temperature. Cell pellets were washed once with GCB and solubilized in 2% Zwittergent 3,14 (EMD Millipore) in PBS for one hour at 37°C. The solubilized suspension was centrifuged for 10 minutes at 2,0000 x g, the supernatant mixed with protein A/G resin (ExAlpha Biologicals) for two hours at 4°C with mixing, and the resin washed three times with 0.5% Zwittergent 3,14 in PBS, and once with PBS alone.

The resin was suspended in 50 µL Laemmli sample buffer without β-mercaptoethanol and subjected to SDS-PAGE for Western blotting; a duplicate gel was run in parallel and stained with Coomassie G-250. Bands from the stained gel were submitted to the Michael Hooker Proteomics Center at the University of North Carolina at Chapel Hill for trypsin digest and identification using mass spectrometry. Accession ID numbers of proteins described in this report are disclosed in Table 1 and Supp. Table 1. Alignment of amino acid sequences was performed using ClustalW.

### Bactericidal assay

A modification of a previously described bactericidal assay [89] was used to test the bactericidal activity of serum from immunized mice. Pooled sera from immunized and control mice were heated at 56°C for 30 min and serially diluted 1:2 in minimal essential medium (MEM) (1:30 – 1:960). Fifty microliters of each dilution were pipetted into wells of a 96-well microtiter plate. Fifty microliters of an MEM suspension containing 100-400 CFU of the target strain were added to the wells and to a well containing 50 µl of MEM alone. After 5 minutes incubation at RT, 50 µl of pooled normal human serum (NHS) (PelFreeze) were added to each well (final concentration 10%) and the plate was incubated for 55 minutes at 37°C in 5% CO_2_. Fifty microliters of GC broth were then added, mixed, and 50 µl aliquots were cultured in duplicate on GC agar and incubated overnight. The antiserum dilution that gave 50% recovery compared to wells without antiserum was defined as the bactericidal_50_ titer. Wells containing heat-inactivated NHS were tested in parallel to measure complement-independent loss of bacterial viability during the assay; no appreciable loss was detected in any experiment. The assay was performed against each strain in two or three independent experiments.

### Statistical analysis

ELISA titers were compared by a Kruskal-Wallis test with Dunn’s multiple comparison. For challenge experiments, the percentage of mice with positive cultures at each time point was plotted for each experimental group as a Kaplan Meier curve and analyzed by the Log Rank test. The number of CFU recovered from vaginal swabs over time was compared by repeated measures ANOVA with Bonferroni correction. The area under the curve (AUC) was calculated for each individual mouse by determining the AUC across the 7 culture time points that was above the limit of detection (20 CFU/mL). Differences between AUC and percentage of vaginal PMNs were compared using a Kruskal-Wallis test with Dunn’s multiple comparison. Statistical analyses were performed using the software Prism (GraphPad Software, La Jolla, CA). Raw data used for statistical analysis of ELISA, *in vivo* efficacy testing, and bactericidal assays have been published [90].

### Animal ethics statement

All animal experiments were conducted at the Uniformed Services University according to guidelines established by the Association for the Assessment and Accreditation of Laboratory Animal Care using a protocol approved by the University’s Institutional Animal Care and Use Committee.

## Acknowledgements

The authors wish to thank Thomas Hiltke, Carolyn Deal and Leah Vincent for helpful discussions, Ian Feavers and Carolyn Vipond for technical advice on ELISAs using 4CMenB as the coating antigen, Melissa Samo for conducting the ELISAs early in this study, and James E. Anderson for providing *Ng* mutant strains, protocols and helpful suggestions. We are also grateful to Rachel Rowland for preparation of media and assistance with animal handling. Mass spectrometry services were provided by the Michael Hooker Proteomics core facility at the University of North Carolina at Chapel Hill.

## Supporting Information

**S1 Fig. Pilot dose response immunization study with the 4CMenB vaccine**. Groups of 5 BALB/c mice were given 20, 100 or 250 µl of the formulated vaccine on days 1 and 28 by the IP or SC routes. (A, B) Serum IgG1 and IgG2a titers against the formulated 4CMenB vaccine 10 days after the second immunization. A dose response is shown for serum IgG1 in both IP- and SC-immunized mice. (C) Serum reactivity against whole cell lysates of *Nm* strain MC58 and of 6 different *Ng* strains using anti-mouse IgG secondary antibody shows a similar dose response based on differences band intensity. A nonparametric test, the Kruskal Wallis test with Dunn’s multiple comparison, was used to analyze ELISA data due to the low sample size. **, p < 0.01.

**S2 Fig. 4CMenB significantly accelerated *Ng* clearance and reduced the *Ng* colonization load in two independent experiments**. In each experiment, mice were immunized three weeks apart with 250-µl doses of 4CMenB by the IP (blue) or SC (red) route or given PBS (black) or alum ([urple) by the IP route (n = 25 or 20 mice per group in experiments 1 and 2, respectively). Three weeks after the final immunization, mice in the diestrus stage or anestrus were treated with 17β-estradiol and antibiotics and challenged with *Ng* strain F62 as described in the Materials and Methods. (A, B) Percentage of culture-positive mice over time and average CFU per ml of a single vaginal swab suspension, respectively for experiment 1 (n = 20-23 mice/group); (C, D) Percentage of culture-positive mice over time and average CFU per ml of a single vaginal swab suspension, respectively for experiment 2 (n = 18-19 mice/group). *, p < 0.05, **p < 0.01, *** p < 0.0001.

**S3 Fig. A similar vaginal PMN influx occurred in all experimental groups**. Vaginal smears collected on each culture day following bacterial challenge were stained with Hemacolor Stain (Sigma), and the percent of PMNs among 100 vaginal cells was determined by cytological differentiation using a light microscopy. An increase in the percentage of PMNs occurred between days 4-7 as is characteristic of this model, with no statistical difference between the groups. The median percent PMNs is shown for each time-point.

**S1 Table. Amino acid identity between proteins of *N. meningitidis* MC58 and *N. gonorrhoeae* FA1090**

